# Mapping whole brain effects of infrared neural stimulation with positron emission tomography

**DOI:** 10.1101/2022.12.24.521746

**Authors:** Marcello Meneghetti, Frederik Gudmundsen, Naja S. Jessen, Kunyang Sui, Christina Baun, Mikael Palner, Christos Markos

## Abstract

The combination of neuroimaging and targeted neuromodulation is a crucial tool to gain a deeper understanding of neural networks at a circuit level. Infrared neurostimulation (INS) is a promising optical modality that allows to evoke neuronal activity with high spatial resolution without need for the introduction of exogenous substances in the brain. Here, we report the use of whole-brain functional [^18^F]fluorodeoxyglucose positron emission tomography (FDG-PET) imaging during INS in the dorsal striatum, performed using a multifunctional soft neural probe. We demonstrate the possibility to identify multi-circuit connection patterns in both cortical and subcortical brain regions within a single scan. By using a bolus plus infusion FDG-PET scanning protocol, we were able to observe the metabolic rate evolution in these regions during the experiments and correlate its variation with the onset of the INS stimulus. Due to the focality of INS and the large amount of viable molecular targets for PET, this novel approach to simultaneous imaging and stimulation is highly versatile. This pilot study can pave the way to further understand the brain connectivity on a global scale.

## 1. Introduction

Understanding the neural functional networks is a crucial goal for the advance of neuroscience and for the development of new treatments for brain diseases [1]. A novel tool towards this goal is the combination of neuromodulation techniques with whole-brain functional neuroimaging to characterize regional activity both locally and distally [2]. Several studies have been already produced combining invasive modulation techniques such as deep brain stimulation (DBS), optogenetic stimulation and chemogenetics, as well as non-invasive ones like focused ultrasound stimulation (FUS) and transcranial direct current stimulation (TDCS), with functional magnetic resonance imaging (fMRI) and positron emission tomography (PET), showing that this approach is highly promising [3, 4, 5, 6, 7, 8, 9, 10, 11, 12, 13]. However, within the invasive techniques, lack of cell-type and spatial specificity, together with the development of scar tissue around chronic implants and complications arising from parasitic Joule heating are well-known issues with electrical stimulation [14]. Optogenetics and chemogenetics excel in cell-type specificity, but they suffer from intrinsic added complexity given by the requirement of genetic manipulation of neurons by either viral injection or use of genetically modified animals. Therefore, their translation to clinical use is limited. Furthermore, the short light wavelengths typically used in optogenetics can generate Becquerel-effect-induced photoelectric artifacts when combined with electrophysiology [15, 16, 17]. Due to these reasons, infrared neural stimulation (INS) has recently gained traction as an alternative focal stimulation technique which can be used both as a non-invasive technique at superficial regions, or through small optic fiber implant with high spatial resolution[18]. This further enables deep structural targeting and simultaneous electrophysiological measurements [19]. The underlying mechanism of INS relies on thermally mediated biological processes that are directly induced by the absorption of infrared (IR) light by biological tissues, and as such no introduction of exogenous substances in the brain is required [20]. Moreover, by varying the operational wavelength within the water absorption spectrum, the penetration length of the stimulation light can be tuned, allowing for spatial resolution down to the individual axon scale[21, 22]. In recent years, the combination of INS and fMRI has been proven to be a powerful tool to study the whole brain functional connectome, with distinct advantages in terms of speed and focality [23, 24].

Between neuroimaging techniques, PET has an unique strength in terms of breadth of applications, as the increasing diversity of radiotracers allows for a wide number of imageable targets and types of measurable outcomes [25]. The use of [^18^F]fluorodeoxyglucose as a radiotracer (FDG-PET), in particular, is the most direct technique to image cerebral metabolism of glucose in the brain, which directly reflects neuronal and glial activity [26, 27]. The introduction of microPET scanners for small animals further increased the sensitivity and accuracy of FDG-PET as an investigation tool [28]. Furthermore, by measuring the brain FDG uptake following a period where the animals is awake and behaving, rather than during the scan, PET measurements can effectively provide information on the brain metabolism in awake small animals, where fMRI is limited by the use of anesthesia or animal restraining during the scan [9]. For applications in combination with INS, FDG-PET also has a distinct advantage over blood oxygenation level dependent (BOLD) fMRI, as resonance frequency shifts due to INS-induced temperature rising might generate non-neural components in the BOLD signal at the stimulation site [24, 11]. Moreover, FDG-PET is not hindered by the presence of metallic implants [29]. This is crucial in enabling precise experiments combining whole brain imaging with local measurements of neuronal activity through electrophysiology during INS.

Electrophysiology is a well-konwn and technologically mature technique for studying neural activity with high sensitivity, temporal resolution and signal-to-noise ratio (SNR), and shares with INS the strength of not needing viral or chemical injections of ‘translators’ [30]. The combination of INS and electrophysiology in vivo in both acute and chronic experimental settings is further facilitated by the recent development of multifunctional implantable neural interfaces able to simultaneously deliver IR light in subcortical brain regions and record electrical signals [31, 32]. Because of all these factors, the combination of INS and PET is a powerful tool in neuroscience, allowing a broad range of complex *in vivo* experiments for the study of brain function in behaving animals on a scale ranging from the single axon to the whole brain.

In this proof of principle study, we combine whole brain dynamic FDG-PET with INS stimulation in the dorsal striatum (DS), similar to our previous target using chemogenetics [11]. We choose this target, as a study case to demonstrate for the first time the combined use of INS and PET for mapping neural circuits *in vivo* (Fig. 1). This allowed us to benchmark the effect of the stimulation on the connectivity within the cortico-striatal-thalamic-cortical (CSTC) circuit, which has been widely studied due to its major involvement in motor disorders such as Parkinson’s disease and Tourette syndrome [33, 34, 35] and behavioural disorders such as obsessive-compulsive disorder and attention deficit hyperactivity disorder [36, 37]. To provide temporal correlation between the stimulation and the metabolism variation, the tracer was slowly infused during the scans to add a temporal dimension to the PET results. The INS stimulus has been delivered using an advanced monolithic fiber interface capable of simultaneous optical stimulation and electrical recording.

**Figure 1:**
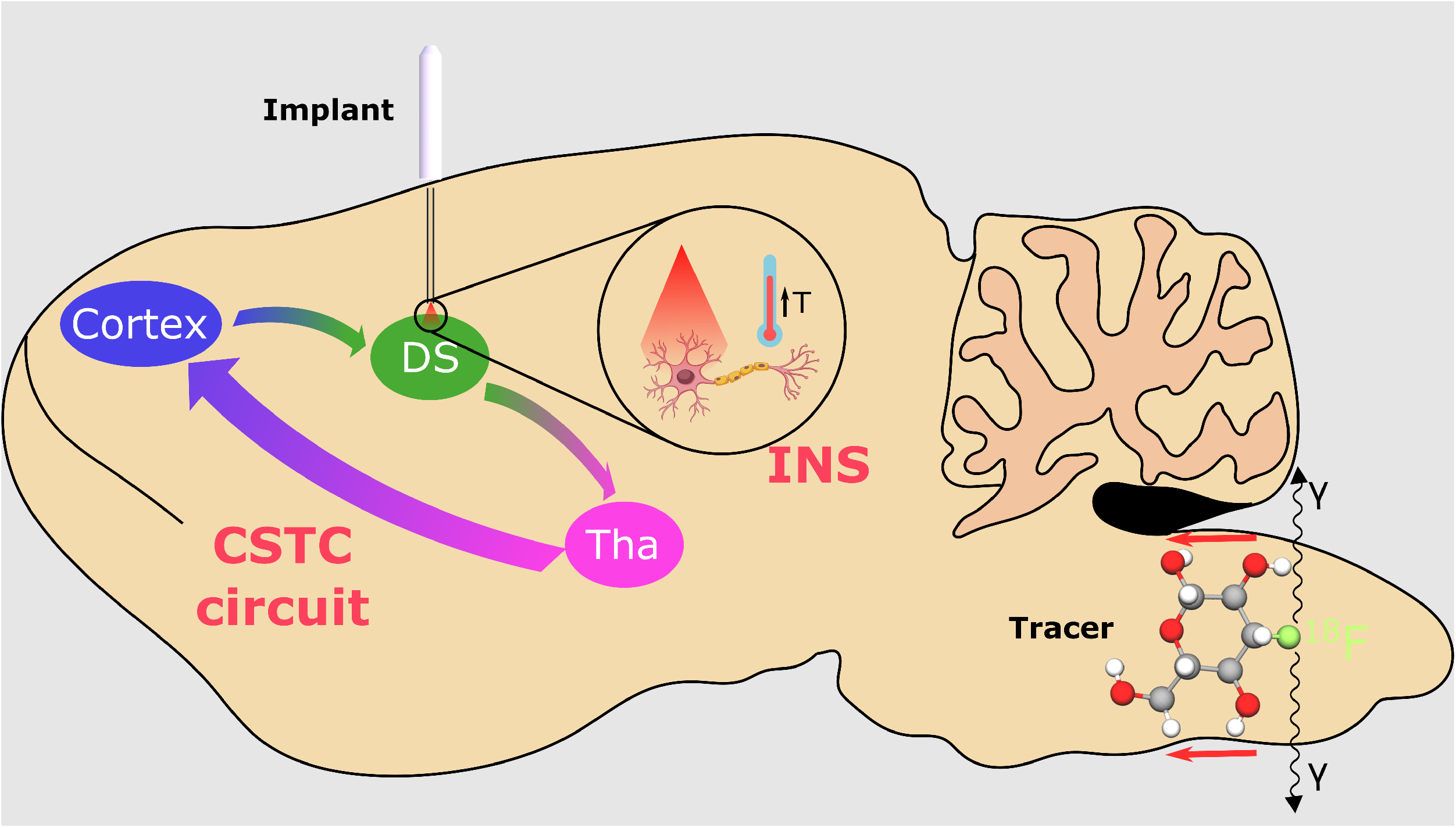
Schematic representation of the study: infrared neurostimulation has been applied to the dorsal striatum using a multifuntional neural interface during the infusion of FDG, and its effects on the CSTC circuits have been evaluated by PET. Tha: thalamus, DS: dorsal striatum

## 2. Materials and Methods

### 2.1. Neural implants

In this study, we used soft infrared neural implants developed by our group for chronic INS applications described in [32] (Fig. 2A). These implants, based on a polymer optical fiber (POF) with a core diameter of ∼ 105 µm, were fabricated by thermal drawing process starting from commercially available polymer rods (Goodfellow UK). Polysulfone (PSU) was used as a core material to achieve enhanced IR light transmission up to 2100 nm wavelength (Fig. 2B,C), while fluorinated ethylene propylene (FEP) was used as a cladding. The use of FEP, one of the softest existing thermoplastics, as a cladding strongly reduced the bending stiffness of the probes with respect to standard silica glass fibers, enhancing their long-term biocompatibility (Fig. 2D) [38]. Hollow microchannels on the side of the core allow for the integration by liquid metal injection of indium electrodes, to enable electrophysiological recording at the fiber tip. The electrodes have an impedance of *<* 35 kΩ in the 1-10000 Hz range (the impedance spectrum of one of the electrodes is shown in Fig. 2E), and have been proven to be suitable for *in vivo* extracellular electrophysiology during INS (Fig. 3). The implants are connectorized with a standard optical fiber ferrule (1.25 mm LC/PC). The implantable length of the interfaces was set at 7 mm for the DS.

**Figure 2:**
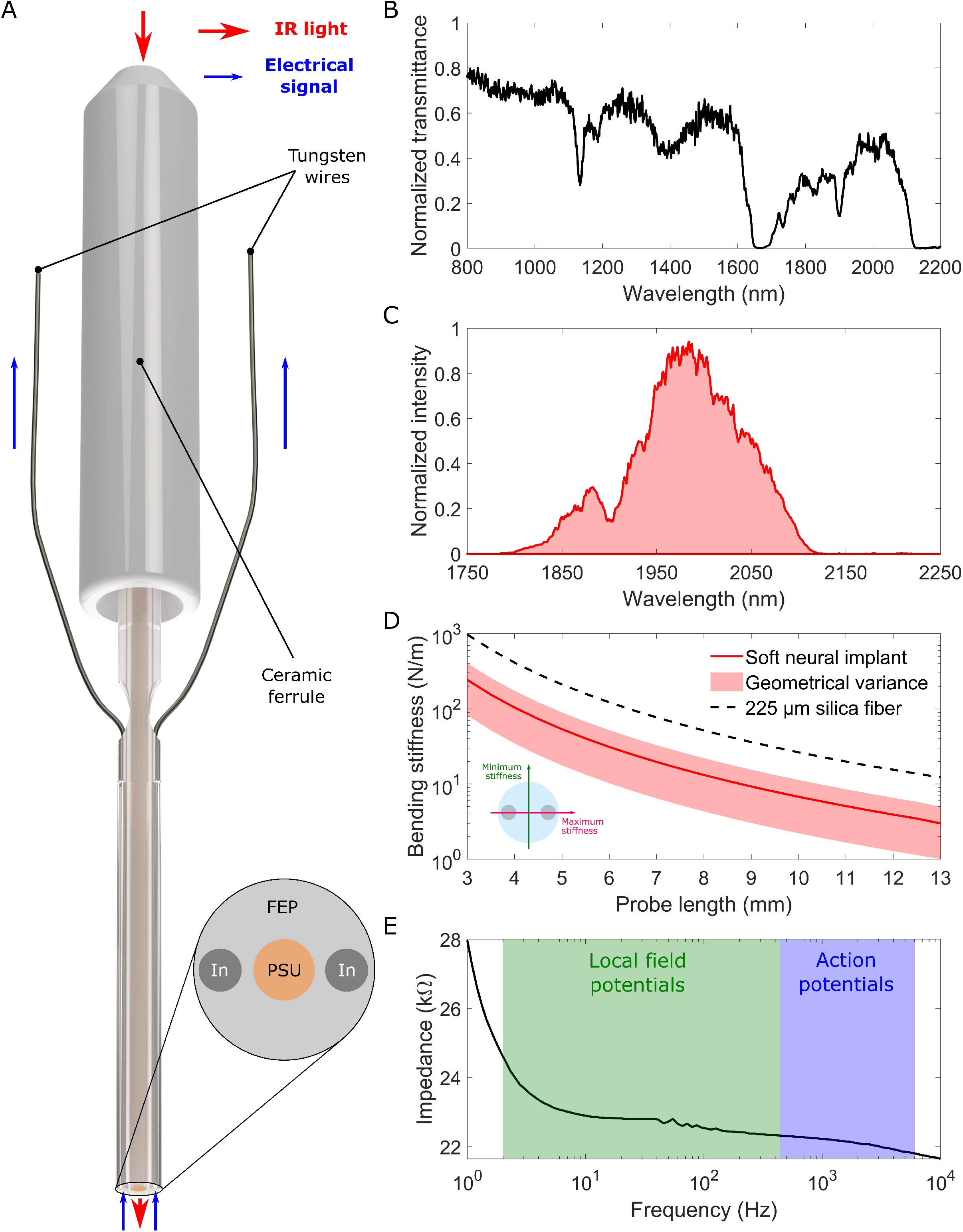
(**A**) Schematic representation of an electrophysiology-eneabled neural interface based on a polymer optical fiber (POF) with a polysulfone (PSU) core, fluorinated ethylene propylene (FEP) cladding functionalized with indium electrodes. (**B**) Normalized infrared transmittance spectrum of the POF. (**C**) Typical impedance spectrum of one of the indium electrodes integrated in our probes. (**D**) Comparison between the average bending stiffness (solid line) of our custom soft neural implant and the one of a standard 225 *µm* silica fiber; the shaded area around the soft probe stiffness represents the minimum and maximum values corresponding to the two main axes of the fiber. (**E**) Normalized output light spectrum of a neural interface connected to the INM setup.

**Figure 3:**
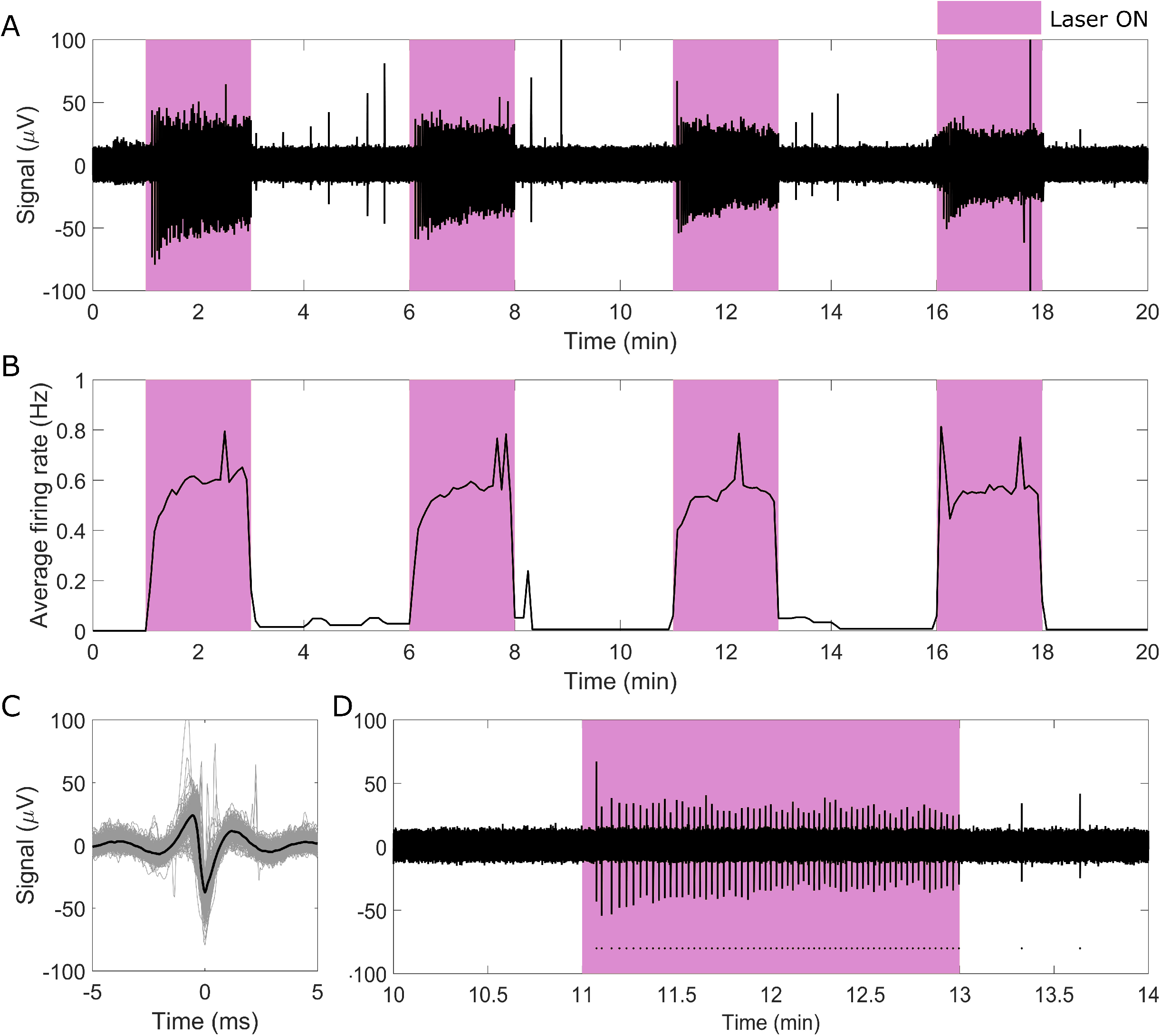
(**A**) Electrophysiological recording performed over several cycles of INS (2 min ON/3 min OFF stimulation). (**B**) Average firing rate measured during the stimulation (5 second bins). (**C**) Overlap (grey) and average (black) of all the recorded spikes. (**D**) Zoom of a single stimulation period, with dots highliting the temporal location of individual spikes

### 2.2. Animals

Six male wild-type adult Long Evans rats (300 ± 50 g for rats in the INS group, 335 ± 13 g for rats in the baseline group) were used for the *in vivo* experimentation. Three of the animals underwent stereotaxic surgery for probe implantation before the PET/CT scans and were used for the INS experiments (INS group), while the other three underwent the PET/CT scans using the same protocol but without prior surgery or stimulation to act as a control group (baseline group). All the procedures performed above have been approved by the Animal Experiments Inspectorate under the Danish Ministry of Food, Agriculture, and Fisheries, and in compliance with the European guidelines for the care and use of laboratory animals, EU directive 2010/63/EU.

### 2.3. Stereotaxic surgery

The animals were all fasted overnight before the scans to ensure low blood sugar and maximal uptake of [^18^*F*]FDG. During all the procedures, anesthesia was maintained using 2% isoflurane in 100% oxygen. Before performing the incision, the head was shaved and, after ensuring the absence of toe pinch response, lidocaine was administered subcutaneously at the site of incision, and ocryl gel was applied to keep the eye moisture during the procedure. Then, the head of the animal was mounted into the stereotaxic frame, where a heating pad connected to an animal temperature controller system was placed below the animal to maintain the body temperature throughout the surgery. A median incision was performed in the scalp to expose the skull. After gently clearing the connective tissue from the surface, the skull was thoroughly washed by successive applications of hydrogen peroxide and saline, and dried with a cotton swab. A small hole (1 mm diameter) was drilled at the target location using an electric dental drill. To avoid any damage to the brain, the dura layer was carefully removed with fine forceps after the drilling. Once the brain tissue was exposed, the neural implant was fixed to the arm of the instrument and slowly inserted into the brain. Finally, the implant was fixed in place with dental cement to prevent any probe movement during the scanning. The stereotaxic coordinates used for the implantations were the following: anterioposterior 1.1mm, mediolateral 2mm, dorsoventral 4mm.

### 2.4. Tracer preparation and administration

The [^18^F]FDG used in this work was from the daily preparation for clinical use (Department of Nuclear Medicine, Odense University Hospital).

### 2.5. PET/CT scan protocol

Following the acute implantation of the neural implant, a catheter was placed in the tail vein of the rat. The animal was then moved to the Siemens Inveon PET/SPECT/CT scanner operating in docked mode (Siemens, Knoxville, Tennessee, USA). They were placed on a heated bed and kept under light isoflurane anesthesia (1.5-2% isoflurane in 100% oxygen). The fiber optic ferrule was connected to the laser setup by means of a silica patch cable and ceramic sleeve, as detailed in the section below. Anatomical CT scans were performed of the head, with the following settings: full rotation in 270-degree projections, 4×4 bin, and low magnification; scan length at 90 mm with an effective pixel size at 111.25 µm and exposure settings 80 kV and 500µA in 350ms. The CT scan was reconstructed using the Feldkamp algorithm with a Sheep-Logan filter, slight noise reduction, and Hounsfield calibration. The animal bed was then extended through the CT unit to make the tail catheter available for infusion. A infusion pump was equipped with a 1mL syringe containing 140-145 MBq [^18^F]FDG in 0.64 mL. A dynamic PET acquisition was started concurrently with the start of [^18^F]FDG infusion (8uL/min). The PET scan was performed in the energy window 350-650 keV and axial scan length of 127mm. Following 40 minutes of baseline recording, INS stimulation was initiated as described below and continued for 20 minutes. Following INS stimulation another 20 minutes of dynamic PET acquisition was collected for a total of 80 minutes uninterrupted acquisition time. The data was histogrammed into 1 minute time frames, starting from 20 minutes, giving a total of 60 frames. CT and PET images were co-registered using a transformation. Reconstruction of PET data was performed using an OSEM3D/SP-MAP algorithm (2 x OSEM iterations and 18 x MAP iterations) with scatter correction and a matrix size 128×128, resulting in a final target resolution of 1.5 mm.

### 2.6. INS protocol

For the INS stimulus, light from a supercontinuum source (NKT SuperK, average power up to 6W) was coupled to a 105 µm core silica patch cable after long-pass filtering with a cut-off wavelength of 1800 nm. Each of the probes was then individually connected to the patch cable using a ceramic sleeve and its output power at different laser currents was measured using a Thorlabs S401C thermal power sensor. This calibration allowed us to account for any small incongruities in light coupling efficiency between the hand-crafted devices. The spectrum at the fiber output was also measured, using an SP320 scanning spectrometer from Instrument Systems (Fig.2.E). Following implantation in the brain, the neural interfaces were reconnected to the patch cable after having placed the animals in the scanner. During the stimulation, light was delivered continuously for 2 minutes intervals, followed by 2 minutes resting periods.

### 2.7. Image and data analysis

Decay-corrected FDG-PET images were analysed in PMOD version 4.2 (PMOD Technologies LLC, Zürich, Switzerland). They were cropped to contain only the head of the animal and matched using an automated pipeline to the Px Rat (W. Schiffer)-FDG-PET template (PMOD v4.2)[39]. Here individual time activity curves from each region of the atlas were extracted and calculated as metabolic uptake in the region divided by whole brain uptake. Each individual time activity curve was then normalized to the baseline metabolic uptake rate during the first 20 minutes, and changes in uptake in relation to baseline plotted as a function of time.

## 3. Results

### 3.1. Targeting during implantation

When using flexible neural interfaces as probes, correctly targeting the selected brain region during implantation without the use of a guide fixture can be a challenge [40]. The importance of the positioning of the probe is even more relevant when using a highly localized stimulation such as the one provided by INS in the 2 µm spectral region. Here, a slow and continuous insertion was used to limit the risk of bending as the fiber penetrates the tissue. The correspondence between the fiber tip location after implantation and the desired stereotaxic coordinates was verified by localizing the fiber in an overlap of PET and CT scans prior to the beginning of the stimulation (Fig. 4). As shown in Figs 4A-C, no significant bending of the fiber with respect to the ceramic ferrule is apparent after insertion. The horizontal slices shown in Figs 4 D-H show the path of the probe through the brain and confirm its location in the targeted region of the right DS.

**Figure 4:**
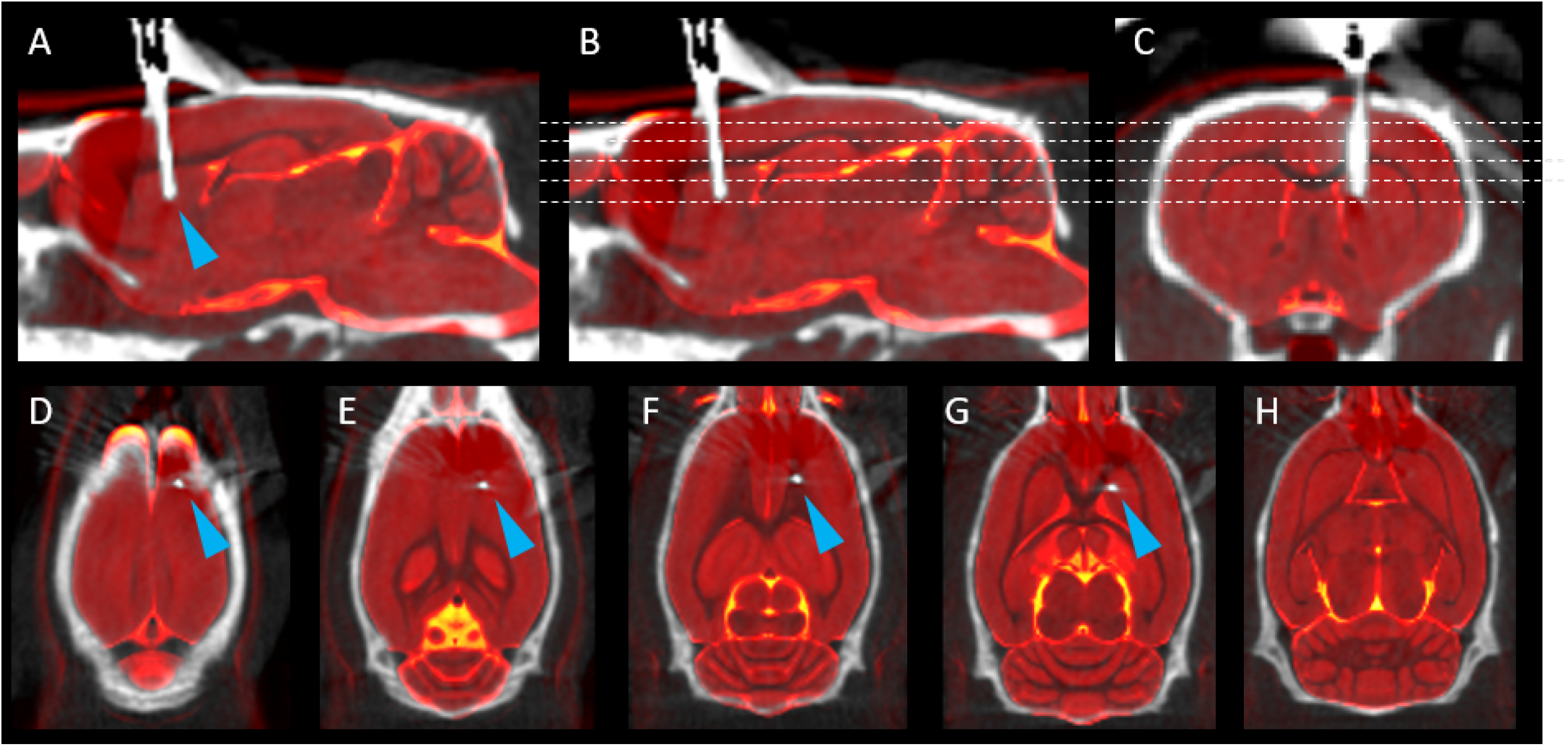
(**A-C**) Sagittal and coronal CT slices (greyscale) overlayed with a standard high resolution structural MR (redscale) showing the location of the neural interface (blue arrow) in the right DS. (**D-H**) Horizontal slices from the planes shown as dashed lines in **B** and **C**, showing the trace of the neural interface through the brain.

### 3.2. Input function

In order to be able to compare the activity variation between different animals, all the metabolic rate results presented in the following sections were normalized to the total signal measured in the animal’s whole brain. Figure 5 shows the input function, i.e. the evolution of this whole brain signal during the radiotracer infusion as a function of the injected dose, for both the INS (A) and baseline (B) groups. While some of the animals slightly deviate from a purely linear behavior of the input function of infusion, this effect is limited to the first minutes of the scans, and as such does not affect the signal recorded for the periods corresponding to the stimulation. This is highlighted by the linear fits of the input function during the 20 to 40 minutes period of the scan (corresponding to the stimulation in the INS group) shown in Figure 5.

**Figure 5:**
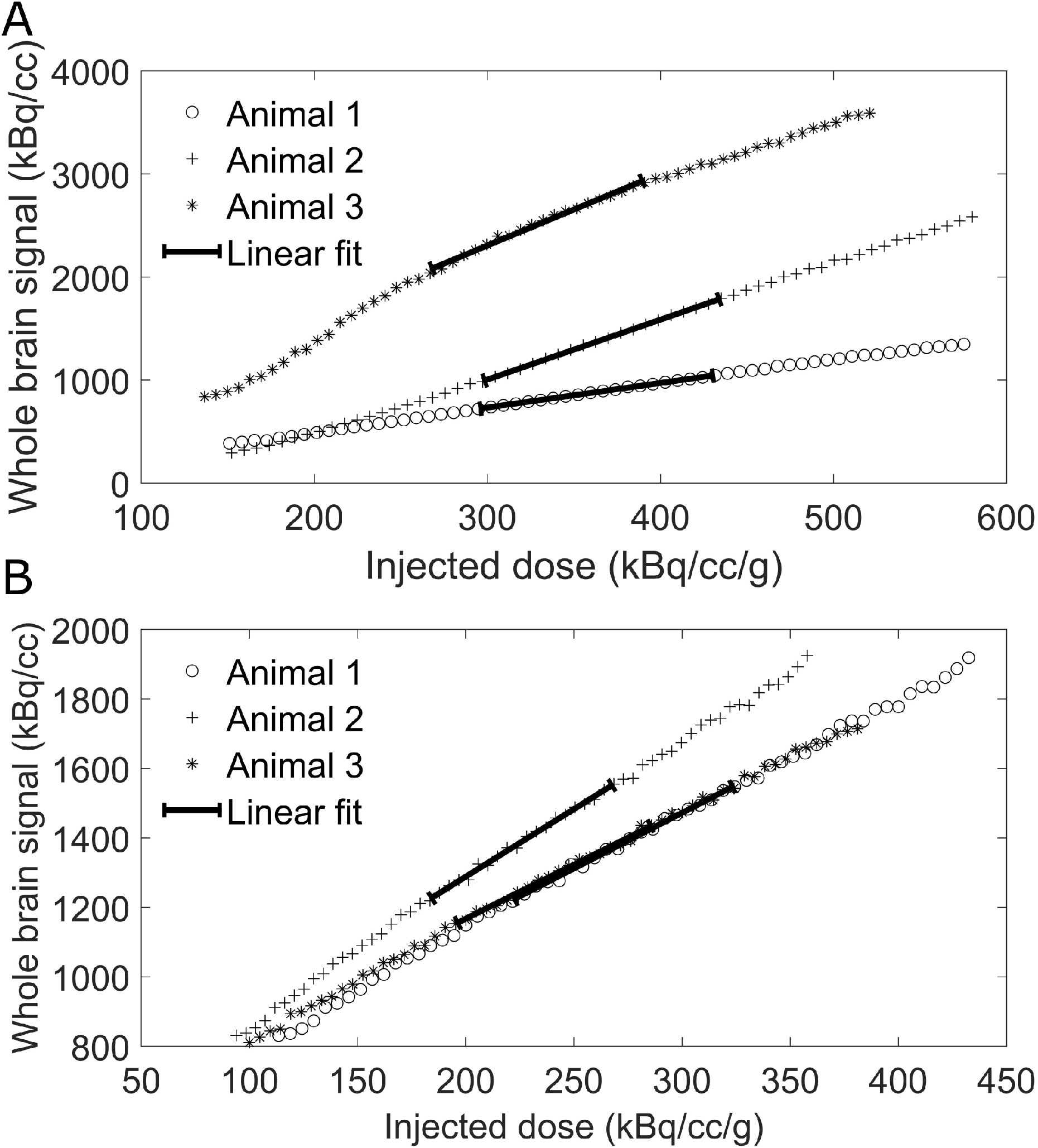
Evolution of the whole brain signal as a function of the injected radiotracer dose for (**A**) the three animals in the INS group and (**B**) the three animals in the baseline group, with linear fits of the data highlighting the period corresponding to the INS stimulus.

The total signal collected from the brain in one of the animals in the INS group is presented in Figure 6A-D, with A presenting the exact location of the neural interface during the scan, and B-D showing all the scans (one per minute) collected before, during and after stimulation, respectively.

**Figure 6:**
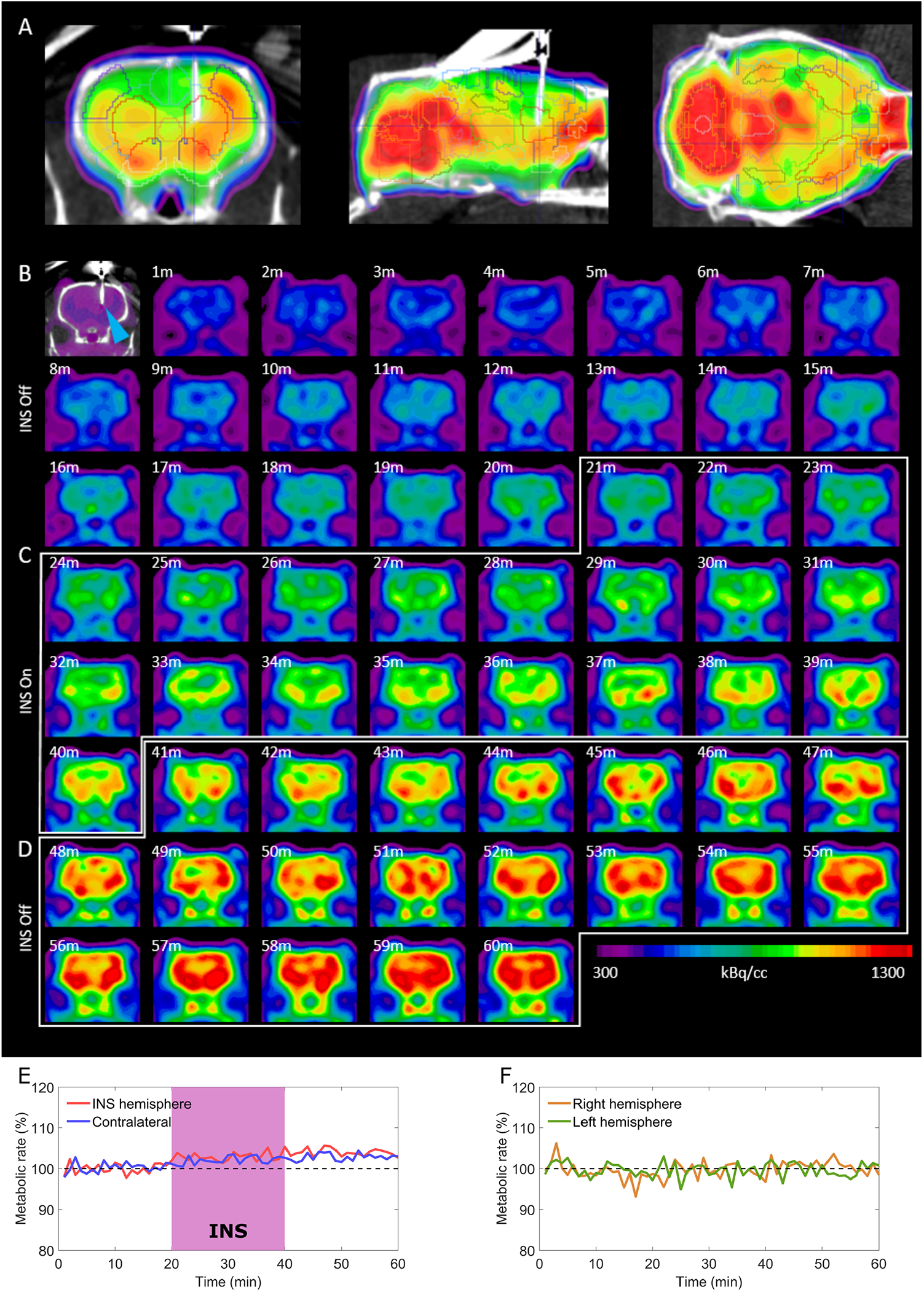
(**A**) Position of the fiber tip in the right DS during the scan, here shown with overlayed volumes of interest. (**B-D**) all frames showing the temporal evolution of the tracer activity before (**B**), during (**C**) and after (**D**) the stimulation. (**E**) Average metabolic rate in the stimulated brain region (striatum) and in its contralateral counterpart before, during and after stimulation (n=3). (**F**) Average metabolic rate in the striatum measured in the animals belonging to the baseline group (n=3).

### 3.3. Metabolic effect of INS on the dorsal striatum

As shown in Fig. 6E, the glucose metabolic rate measured in the DS that was directly targeted by stimulation (right hemisphere) resulted to be visibly higher than the whole brain average for the rats in the INS group. For rats in the baseline group, instead, the metabolic rate of the right DS was aligned to the whole brain average (Fig. 6F). Furthermore, the acceleration of the glucose metabolism is coincident in time to the onset of the stimulation, showing a temporal correlation between the INS and the measured effect which is consistent with the preliminary electrophysiological data presented in Fig.3. These results confirm that the selected INS protocol is effective in initiating neural activity in the region surrounding the fiber tip. After the end of the stimulation, the metabolic rate remains stable but elevated, showing a protracted effect of INS. Consistently with existing literature that depicts a pronounced bilaterality of functional connectivity involving the striatum within circuits in the basal ganglia system [41], we observed a similar metabolic rate variation in the contralateral DS which is present in the INS group animals and not in the baseline group animals (Fig. 6E,F).

### 3.4. Deep brain involvement

According to the well-known model of the CSTC circuit, excitation of the striatum can result in either excitatory or inhibitory stimulation of the thalamus (direct and indirect pathways, respectively) [42]. As presented in Figure 7A, in the animals belonging to the INS group we observed a decrease of the thalamic activity in the hemisphere ipsilateral to the INS location, indicating that the infrared stimulus could be favoring the indirect pathway with respect to the direct one. The reduced ipsilateral midbrain activity shown in Fig. 7B and the increased activity in several cortical regions presented in the next section, however, hint at the involvement of several cortical micro-circuits in addition to the CSTC. The INS-induced variation in the bilateral activity of both hypothalamus and VTA visible in Fig. 7C and D, in particular, seems to indicate the involvement of cortico-striato-hypothalamic limbic circuits involved in feeding behavior, despite their usual association with the ventral part of the striatum rather than the DS [43, 44]. As for the striatal effect described in the previous section, no change of metabolic rate during the scan was observed in the animals in the baseline group. In all the considered regions, the temporal coincidence between the change in metabolic rate and the onset of the stimulus confirms that the observed alterations in brain activity can be attributed to INS.

**Figure 7:**
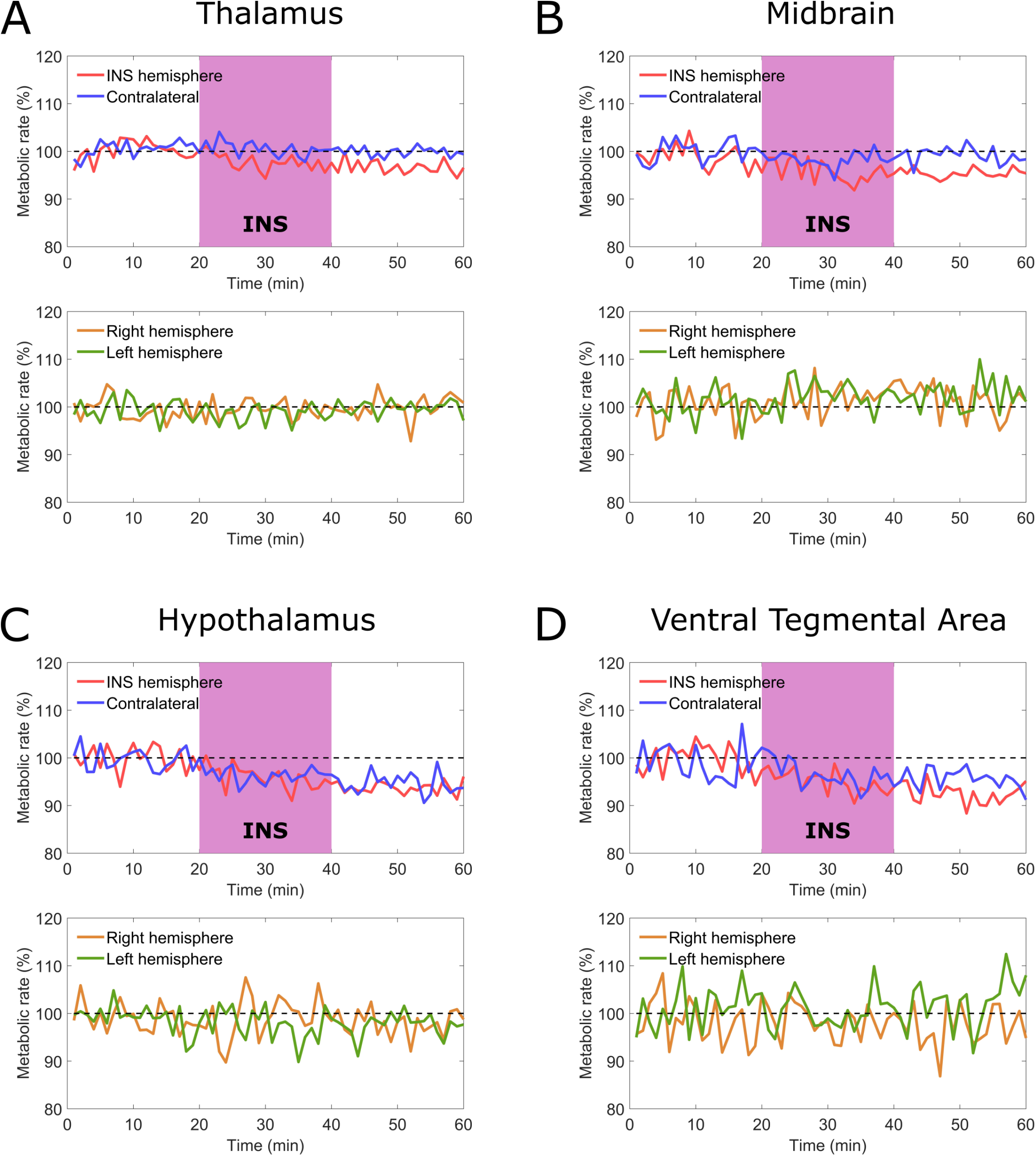
Evolution of the metabolic rate in time in different deep brain regions for the rats in the INS group (n=3, blue and red lines) before, during and after stimulation. The metabolic rate is compared with the one of the rats in the baseline group (n=3, orange and green lines). The considered regions are hypothalamus, midbrain, thalamus and ventral tegmental area

### 3.5. INS induced recruitment of cortical brain regions

While the stimulation of the indirect pathway in the CSTC circuit and the related inhibition of thalamic activity are expected to correspond to an equivalent inhibition of activity in the PFC, the FDG signal measured in the medial PFC was affected by a level of noise too high to be able to discriminate variations in the metabolism of this region between the INS group and the baseline group (Fig. 8A). Despite the measured reduction in the metabolism of the thalamus, we observed a bilateral increase in the activity in the motor cortex for the animals undergoing stimulation, which was particularly pronounced in the hemisphere ipsilateral to the stimulation (Fig. 8B). This cortical region is strongly associated with the classical description of the CSTC circuit involving the dorsal striatum [45]. Interestingly, the INS stimulus also affected the activity of both orbitofrontal and entorhinal cortex (Fig. 8C and D), once again suggesting the involvement of the ventral circuitry described in [44].

**Figure 8:**
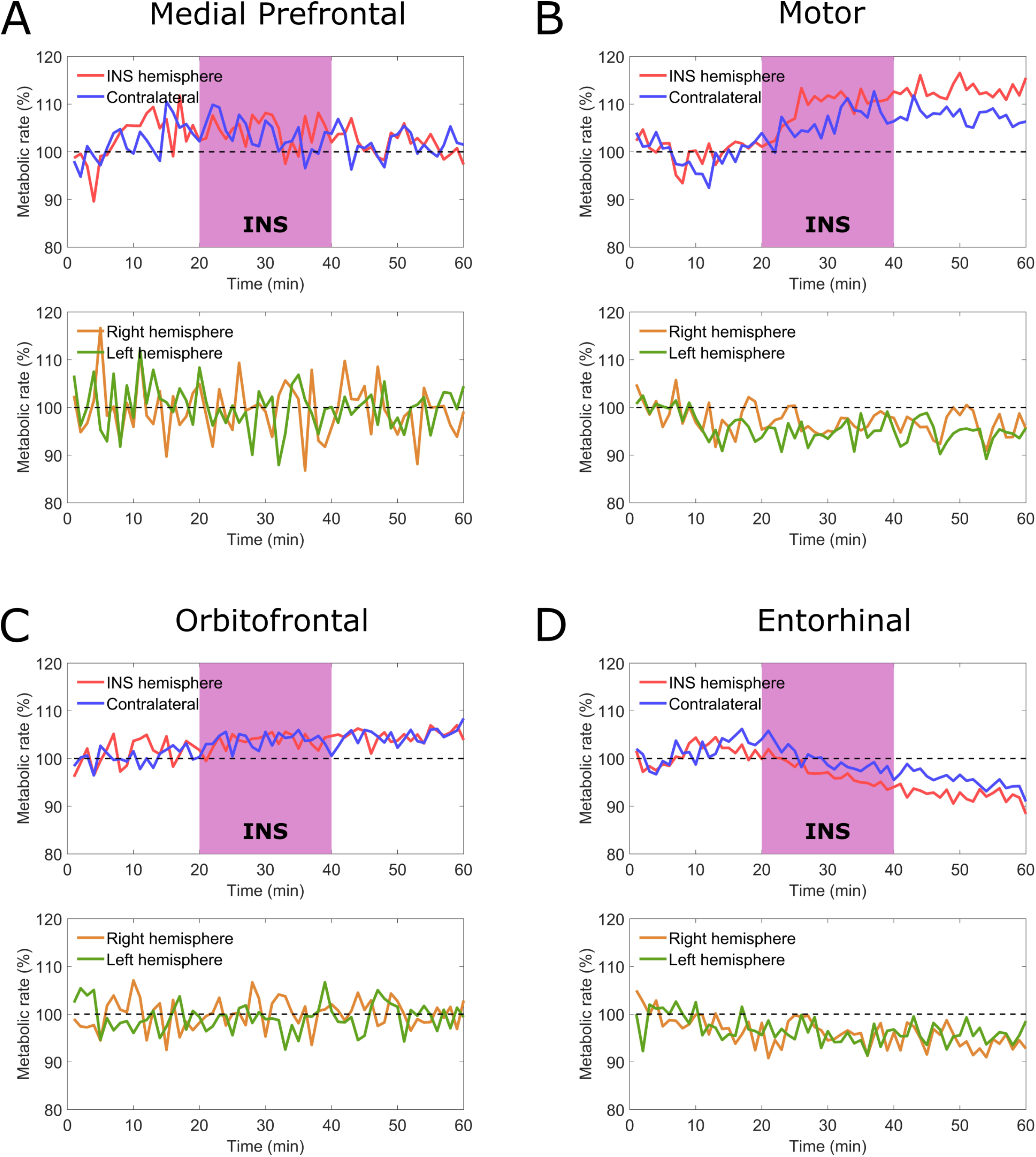
Evolution of the metabolic rate in time in different cortical regions for the rats in the INS group (n=3, blue and red lines) before, during and after stimulation. The metabolic rate is compared with the one of the rats in the baseline group (n=3, orange and green lines). The considered regions are enthorinal cortex, medial prefrontal cortex, motor cortex and orbitofrontal cortex

## 4. Discussion

We demonstrated the feasibility of using a combination of INS and FDGPET for mapping neural functional networks *in vivo* on a global scale. By using broadband supercontinuum IR light combined with custom-made soft bidirectional neural interfaces, we stimulated the dorsal striatum in anesthetized animals to evaluate the functionality of this novel approach on the CSTC circuit. We obtained a temporal profile of the glucose metabolism in different brain regions by scanning at regular (one minute) intervals. This allowed us to temporally correlate the variation of the metabolic rate in all considered regions with the onset of the INS stimulus, thus excluding the possibility of the effects being caused by the implant and eliminating the need for sham experiments. The stimulation protocol, which was already validated by electrophysiological recordings in our previous study [32], was confirmed by the increase of the metabolic rate in the stimulated region. Global scale mapping of the glucose metabolism during the stimulation allowed to observe a rich network of connections in both the ipsi- and contralateral hemispheres. This involvement of regions distal to the stimulated areas spanned from the cortex to the deep brain. Interestingly, this included regions not typically considered as a part of the CSTC circuit, which was the focus of this study. The presence of bilateral variation in the hypothalamic region, as well as in the orbitofrontal and entorhinal cortex suggest the additional involvement of the cortico-striato-hypothalamic circuitry.

In summary, INS and PET can be combined to form a powerful mapping tool for brain circuitry *in vivo*. The high spatial specificity and lack of need for viral manipulation of INS, combined with the versatility of molecular targets for PET, can provide a powerful tool to rapidly study brain connectivity with high precision while decoding the studied networks on a chemical level. Novel infrared interfaces based on soft materials, as the ones used in this study, open possibilities for chronic implantations with minimal inflammation and, subsequently, to studies in behaving animals. We believe this work will pave the way to future investigations in different subcortical brain region and thus represent a cornerstone for further advances in our understanding of the inner mechanisms of the brain.

## Data and code availability statement

All software and procedures concerning the data acquisition and analysis have been detailed in the Material and Methods section. All data sets are available from the corresponding author upon reasonable request.

## Ethics approval statement

All the procedures performed above have been approved by the Animal Experiments Inspectorate under the Danish Ministry of Food, Agriculture, and Fisheries, and in compliance with the European guidelines for the care and use of laboratory animals, EU directive 2010/63/EU.

## Acknowledgments

This research has been financially supported by Lundbeck Fonden projects (Multi-BRAIN, R276-2018-869 and R380-2021-1171) and VILLUM FONDEN (36063).

